# SNPWay: streamlined SNP-to-function and pathway over-representation analysis

**DOI:** 10.64898/2026.05.03.722523

**Authors:** Bryan Queme, Ayaan Kakkar, Anushya Muruganujan, Paul D. Thomas, James W. Gauderman, Huaiyu Mi

## Abstract

**Motivation:** Post-GWAS interpretation frequently requires translating variant lists (e.g., lead SNPs, clumped loci, credible sets, or curated panels) into pathway and functional hypotheses. In practice, obtaining pathway and functional over-representation results from SNP inputs often requires stitching together multiple tools for variant annotation, regulatory annotation, gene identifier handling, and statistical testing. This integration burden can reduce reproducibility and restrict end-to-end analysis to groups with dedicated bioinformatics support.

**Summary:** We present SNPWay, a web server and R package that performs end-to-end SNP-to-function and pathway over-representation analysis in a single standardized workflow. SNPWay accepts rsIDs, VCF files, or hg19/GRCh37 genomic coordinates. It queries Annotation Query (AnnoQ) to obtain SNP-to-gene mappings from ANNOVAR, SnpEff, and VEP under both Ensembl and RefSeq gene models, and incorporates enhancer-gene links via PEREGRINE to augment mappings for noncoding variants. SNPWay aggregates mapped genes into a single, non-redundant, combined gene list and submits it to PANTHER for over-representation testing against the Homo sapiens reference list, returning over-represented pathways and functional categories (e.g., Gene Ontology) with direct links for interactive exploration in PANTHER. SNPWay’s modular architecture is designed for extensibility, enabling incorporation of additional analysis methods in future releases. A step-by-step walkthrough is provided in Supplementary Data.

**Availability and implementation:** Web server: https://snpway.annoq.org/.

R package and source code: https://github.com/USCbiostats/Annoq_Overrepr_Workflow.

Documentation: https://snpway.annoq.org/about.

Examples: The website contains ‘Sample files’ for the input formats, also provided in Supplementary Data. SNPWay is free to use with no mandatory login.

**Supplementary information:** Supplementary data are available at *Bioinformatics* online

## 1 Introduction

Genome-wide association studies (GWAS) have identified large numbers of trait-associated single-nucleotide polymorphisms (SNPs)^1–4^, yet the biological interpretation of these variants remains a central challenge. A common post-GWAS strategy is to map SNPs to genes and then evaluate pathways or gene-set over-representation to identify underlying biological implications^5^. Several platforms support components of this workflow, such as FUMA^6^, which provides modular functionality for SNP-to-gene mapping and downstream functional enrichment, primarily in the context of gene prioritization from GWAS summary statistics.

A critical but often under-appreciated step in this pipeline is SNP-to-gene annotation. Widely used variant annotation tools, including ANNOVAR^7^, SnpEff^8^, and the Ensembl Variant Effect Predictor (VEP)^9^, rely on different algorithms and transcript definitions, and their outputs depend on the underlying gene model, such as Ensembl^10^ and RefSeq^11^. Consequently, SNP-to-gene assignments can differ across commonly used annotation configurations, and these differences can propagate to downstream pathway analyses^12–15^. In addition, because many GWAS signals lie outside coding regions, enhancer-gene relationships can provide regulatory context for associating noncoding variants with plausible target genes^16^.

Although annotation choices can influence variant interpretation^12–15^, most post-GWAS workflows implicitly adopt a single annotation strategy. In contrast, statistical gene-set analysis methods such as MAGMA^17^ focus on robust aggregation of association statistics, but generally assume a fixed SNP-to-gene mapping has already been defined and formatted for downstream testing. As a result, moving from common SNP inputs (rsIDs or VCFs) to pathway over-representation results often requires bespoke scripting and identifier handling, limiting reproducibility and accessibility.

Several tools address part of this problem but differ in scope and design. snpXplorer^18^ provides SNP-based functional annotation and pathway enrichment via g:Profiler^19^, but relies on a single annotation strategy and does not integrate enhancer-gene links. GSA-SNP2^20^ and Pascal^21^ offer pathway-level scoring from GWAS summary statistics but operate on pre-computed SNP p-values rather than accepting variant lists directly. XGR^22^ supports enrichment analysis of SNP and gene lists across multiple ontologies but does not combine annotations from different tools or gene models. FUMA provides comprehensive post-GWAS functionality, including positional, eQTL, and chromatin interaction mapping, but requires full GWAS summary statistics and a mandatory login, and does not aggregate SNP-to-gene assignments across multiple annotation algorithms. To our knowledge, no existing tool provides a single, login-free workflow that (i) accepts common variant-list inputs (rsIDs, coordinates, or VCF files), (ii) aggregates SNP-to-gene mappings from multiple annotators (ANNOVAR, SnpEff, VEP) across both Ensembl and RefSeq gene models, (iii) incorporates enhancer-gene regulatory links, and (iv) submits the combined gene set directly for pathway overrepresentation and gene ontology testing.

To address this gap, we developed SNPWay, a graphical user interface (UI) and R package for SNP-to-function and pathway over-representation analysis in a single standardized workflow. SNPWay leverages AnnoQ API^23^ to retrieve SNP-to-gene annotations from multiple annotators (ANNOVAR, SnpEff, VEP) across both Ensembl and RefSeq gene models, and incorporates enhancer-linked target genes via PEREGRINE^16^, as many GWAS signals are non-coding. SNPWay then submits the combined mapped gene set to PANTHER API^24^ for over-representation^25^ using the *Homo sapiens* reference list. SNPWay returns enrichment results and the combined gene list used for testing, supporting streamlined reporting and reproducible execution through both UI and R package.

## 2 Implementation

SNPWay is delivered as (i) an interactive web interface for on-demand analysis and result download, and (ii) an R package for scripted, reproducible analyses and batch workflows. Both interfaces execute the same pipeline steps and produce the same output schema (Figure 1).

**Figure 1.**
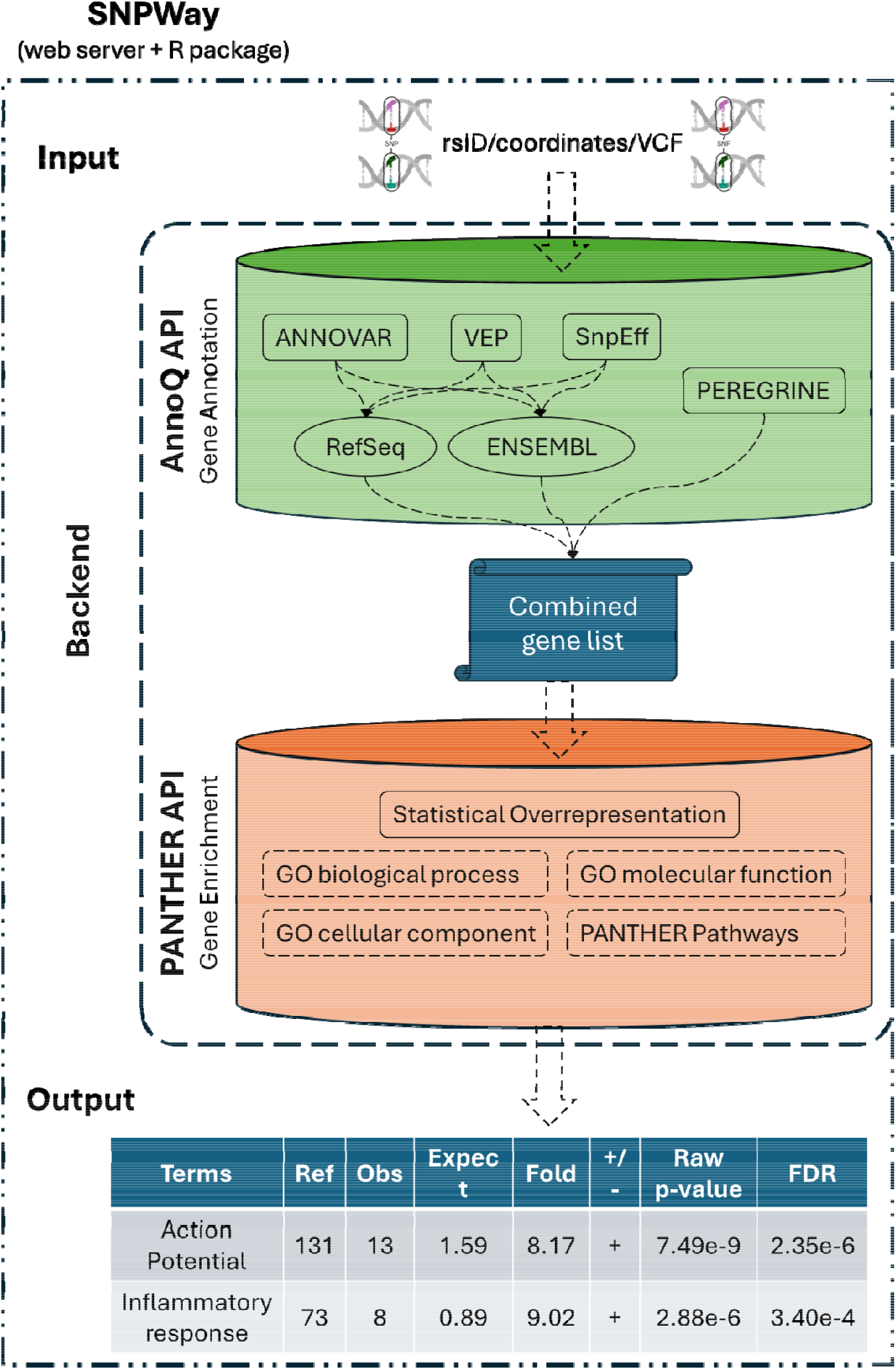
SNPWay workflow. SNPWay accepts rsIDs, hg19/GRCh37 coordinates, or VCF inputs and normalizes variants. SNPWay queries AnnoQ API to retrieve SNP-to-gene mappings from ANNOVAR, SnpEff, and VEP under Ensembl and RefSeq gene models and incorporates enhancer–gene links via PEREGRINE. Unique mapped genes are aggregated into a combined gene set and submitted to PANTHER’s over-representation test using the Homo sapiens reference list. SNPWay returns enrichment results (FDR-adjusted significance, fold enrichment, observed/expected counts, direction) and provides a direct link for interactive exploration in PANTHER.

### 2.1 Data input

SNPWay accepts human hg19/GRCh37 variant input in three common formats: rsIDs, VCF files, or genomic coordinates (chromosome/position). VCF inputs are parsed to extract variant coordinates; only CHROM, and POS are required. The additional identifiers a VCF file may have are optional.

SNPWay is intended for variant-list interpretation (e.g., lead SNPs, curated panels, clumped loci, credible sets) rather than LD-aware association modeling. Users provide a pre-defined variant set appropriate for their study design.

SNPWay currently operates on hg19/GRCh37 coordinates, consistent with the AnnoQ annotation database; a lift-over module to accept hg38/GRCh38 inputs is planned for a future release.

### 2.2 SNP-to-gene mapping via AnnoQ

SNPWay retrieves SNP-to-gene mappings by querying the AnnoQ API, which provides standardized access to ANNOVAR, SnpEff, and VEP annotations. For each input variant, SNPWay collects gene assignments under both Ensembl and RefSeq gene models and incorporates enhancer-linked target genes from PEREGRINE to provide regulatory context for noncoding and intergenic variants. Gene mappings are aggregated as a union across all annotator–model combinations to produce a single combined gene list. Because annotation tools differ in their algorithms^12,14^ and transcript definitions^13^, individual tools can miss relevant gene assignments; prior work has shown concordance as low as 65% for loss-of-function annotations between tools using the same transcript set^12^. Taking the union mitigates these gaps and potentially provides a more comprehensive input for downstream enrichment testing^15^. Importantly, prior benchmarking showed that this union strategy does not inflate gene lists with spurious assignments but rather recovers biologically relevant mappings missed by individual tools^15^. The combined gene list is submitted to PANTHER for over-representation testing.

### 2.3 Functional and pathway over-representation analysis using PANTHER (human reference list)

The detailed step-by-step tutorial can be found at https://snpway.annoq.org/tutorial.

SNPWay submits the final combined gene set to PANTHER for over-representation analysis through Fisher’s exact test with False Discovery Rate (FDR) adjustment by default, using the Homo sapiens reference list.

Additionally, SNPWay supports all functional categories exposed through PANTHER’s workflow^24^ (e.g., Gene Ontology classes and pathway annotations). Returned results include FDR-adjusted significance, fold enrichment, observed and expected counts, and direction (over- or under-represented), consistent with PANTHER outputs.

Because PANTHER performs identifier mapping internally, SNPWay submits the combined gene list without an additional harmonization step; genes that do not map are handled within the PANTHER results interface, which users can access via the “View full results in PANTHER” link.

For each run, SNPWay returns an input normalization report, the combined gene list submitted for testing, and PANTHER over-representation results tables (FDR, fold enrichment, observed/expected counts, direction), with downloadable outputs and a direct link to open full results in PANTHER.

Advanced options also allow users to select between Fisher’s exact test and the binomial test, and to apply FDR, Bonferroni, or no multiple-testing correction (defaults: Fisher’s exact test with FDR). This flexibility allows the same SNP-to-function pipeline to be applied across different functional classification schemes and statistical frameworks depending on the interpretive goals of the analysis.

Results are presented in a PANTHER-style table within SNPWay and can be exported for reporting.

### 2.4 Modular architecture and extensibility

SNPWay is built on a modular architecture in which variant input handling, annotation retrieval, gene aggregation, and enrichment testing are implemented as independent, composable components (Figure 1). Individual pipeline stages can be updated, replaced, or extended without modifying the overall workflow. For example, the annotation retrieval module could be extended to incorporate eQTL-based gene mappings, and the enrichment module could be supplemented with gene-set enrichment analysis or network-based methods. We plan to leverage this design in future releases to support additional analysis types beyond pathway over-representation testing such as pPRS^26^.

### 3 Use case

We demonstrate SNPWay by reproducing the annotation and enrichment steps of a published pathway polygenic risk score (pPRS) workflow applied to colorectal cancer (CRC)^26^ and provide an additional colorectal cancer example application examining red meat intake and a TGF-β pathway–based pPRS^27^. The step-by-step instruction for this use case can be found in the Supplementary Data. In the pPRS study, 204 CRC-associated GWAS SNPs were submitted to SNPWay, which mapped 189 variants to 265 protein-coding genes via the multi-annotator union and PEREGRINE enhancer links. PANTHER over-representation testing identified four significantly enriched pathways (FDR < 0.05), including TGF-β signaling, Alzheimer disease-presenilin, Gonadotropin-releasing hormone receptor, and Cadherin signaling pathway. These pathway-defined variant subsets were then used to construct pPRS that revealed a significant pPRS × NSAIDs interaction (p = 0.0003) in a 78,253-subject CRC study, an interaction not detected using the overall PRS (p = 0.41).

The pPRS workflow above used PANTHER pathway annotations for over-representation testing. However, SNPWay supports the full range of functional categories available through PANTHER, including Gene Ontology (Biological Process, Molecular Function, Cellular Component), PANTHER GO-Slim categories, PANTHER Protein Class, and Reactome pathways (Supplementary Data).

## 4 Conclusion

SNPWay provides an accessible, end-to-end workflow that converts SNP inputs (rsIDs, hg19/GRCh37 coordinates, or VCF files) into functional and pathway over-representation results in a single run. By integrating established variant annotation outputs across multiple tools and gene models and incorporating enhancer– gene links through PEREGRINE, SNPWay lowers the practical barrier to pathway-focused interpretation of GWAS variants. SNPWay returns enrichment tables, supporting transparent reporting and reproducible downstream analyses. Its modular architecture is designed to accommodate future extensions, including alternative enrichment methods and additional regulatory annotation sources, broadening the scope of functional interpretation available from variant-list inputs.

## Acknowledgements

We would like to thank the AnnoQ and PANTHER software engineers Dustin Ebert and Tremayne Mushayahama for working with us to make this software possible. We also would like to thank Mingzhi Ye for his input in the early stage of the development of this tool.

## Author Contributions

Bryan Queme (Conceptualization [equal], Methodology [lead], Software [equal], Validation [lead], Writing – original draft [lead], Writing – review & editing [equal]), Ayaan Kakkar (Software [lead]), Anushya Muruganujan (Software [equal], Visualization [equal]), Paul D. Thomas (Supervision [equal]), James W. Gauderman (Conceptualization [equal]), Huaiyu Mi (Conceptualization [equal], Methodology [equal], Supervision [equal], Writing – review & editing [equal]).

