## Supplemental Data for "SNPWay: streamlined SNP-to-function and pathway over-representation analysis"

We demonstrate SNPWay by reproducing the pathway over-representation analysis from Gauderman et al. using the 204 CRC-associated GWAS SNPs.

1. Go to the pPRS paper: <https://doi.org/10.1371/journal.pgen.1011543>
2. Download S4 Table. Gene and pathway annotation for 204 colorectal-cancer-associated SNPs.
3. Select the 306 rsIDs entries (representing 204 unique SNPs; only unique rsIDs will be used for analysis)
4. In SNPWay, under “Retrieve SNP data by” select rsIDs. The user can also use the VCF File and Chromosome position options.

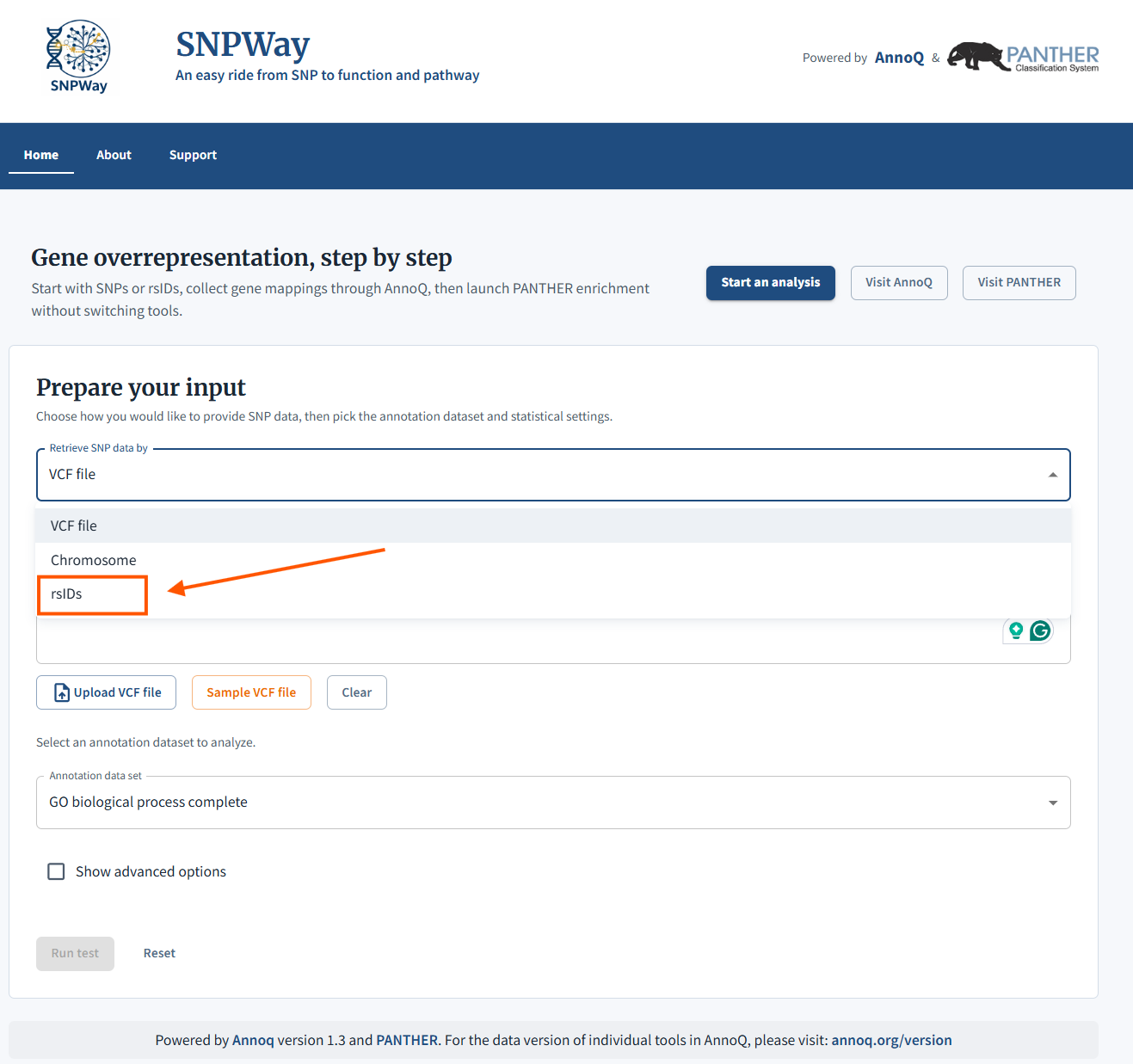

5. Paste the rsIDs from the pPRS paper

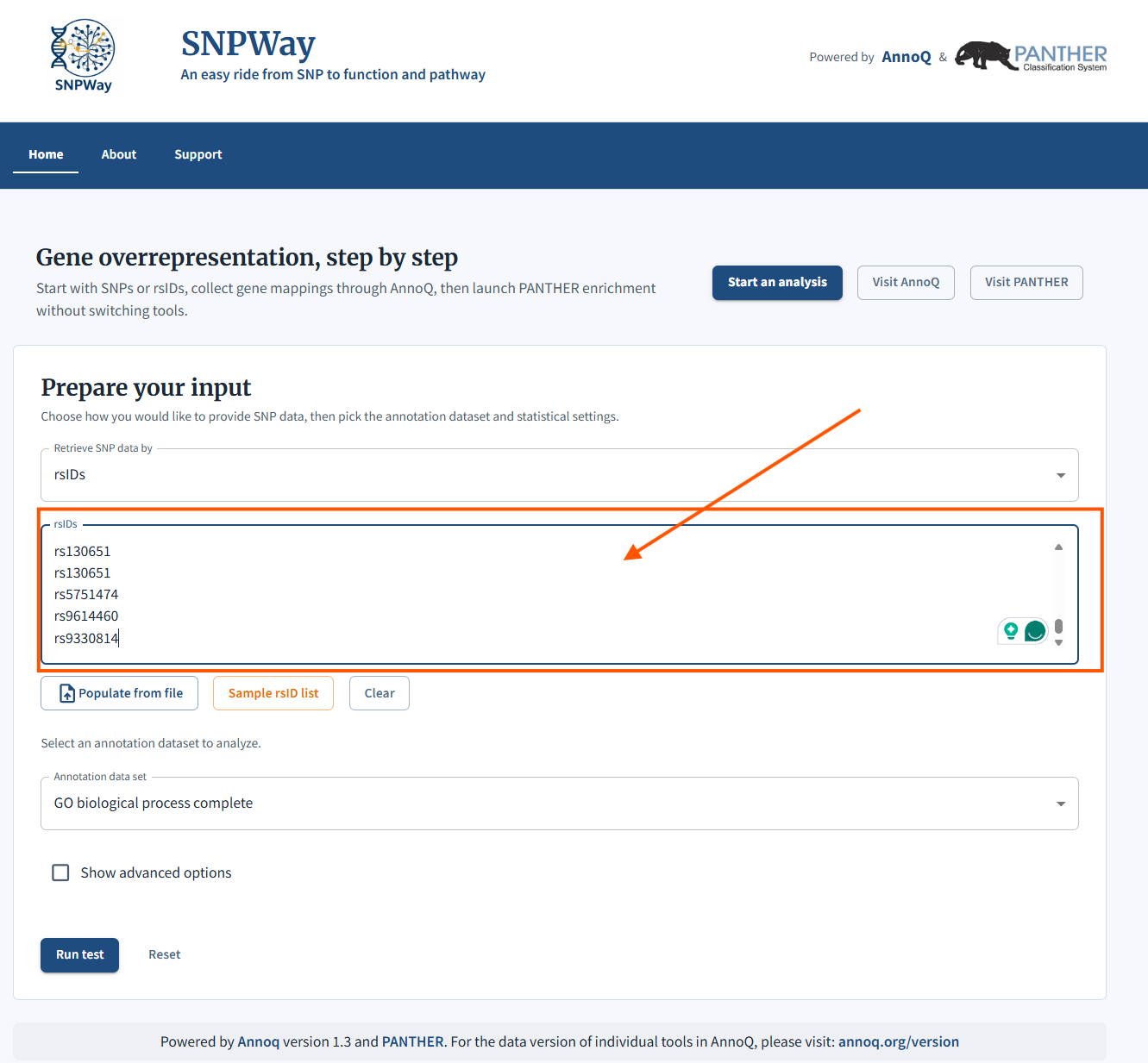

6. Select from the “Annotation data set”.
   a. For the pPRS, we will select “PANTHER pathways”. For this walkthrough we select PANTHER pathways; other available options include Gene Ontology (Biological Process, Molecular Function, Cellular Component), PANTHER GO-Slim categories, PANTHER Protein Class, and Reactome pathways. Users can repeat this workflow with any annotation dataset to explore different functional interpretations of the same variant set.

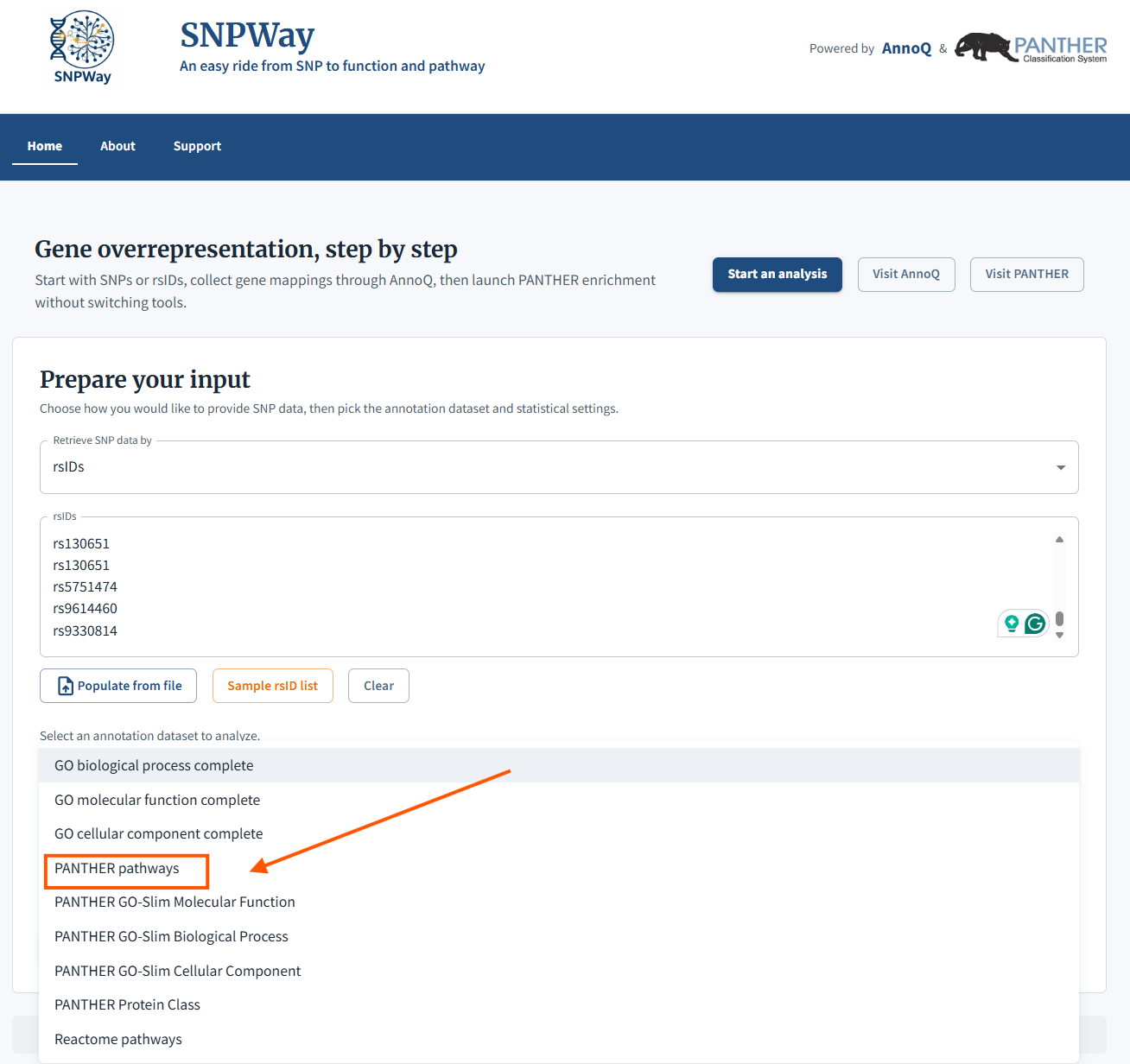

7. From the advanced options, you will be able to choose the “Test Type” and “Correction”. The default is Fisher’s exact with FDR correction

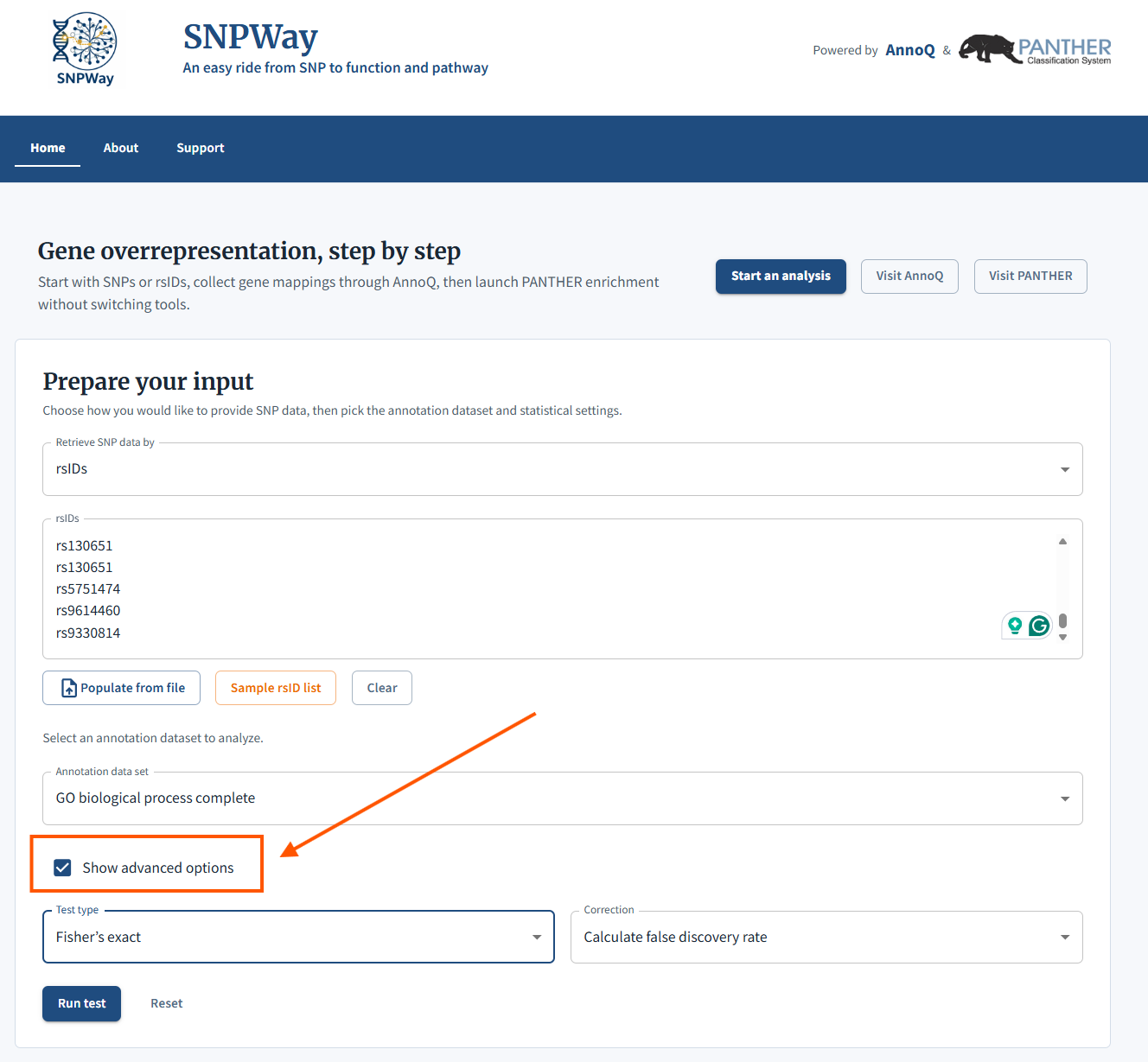

8. Lastly, we will “Run test” to submit the job

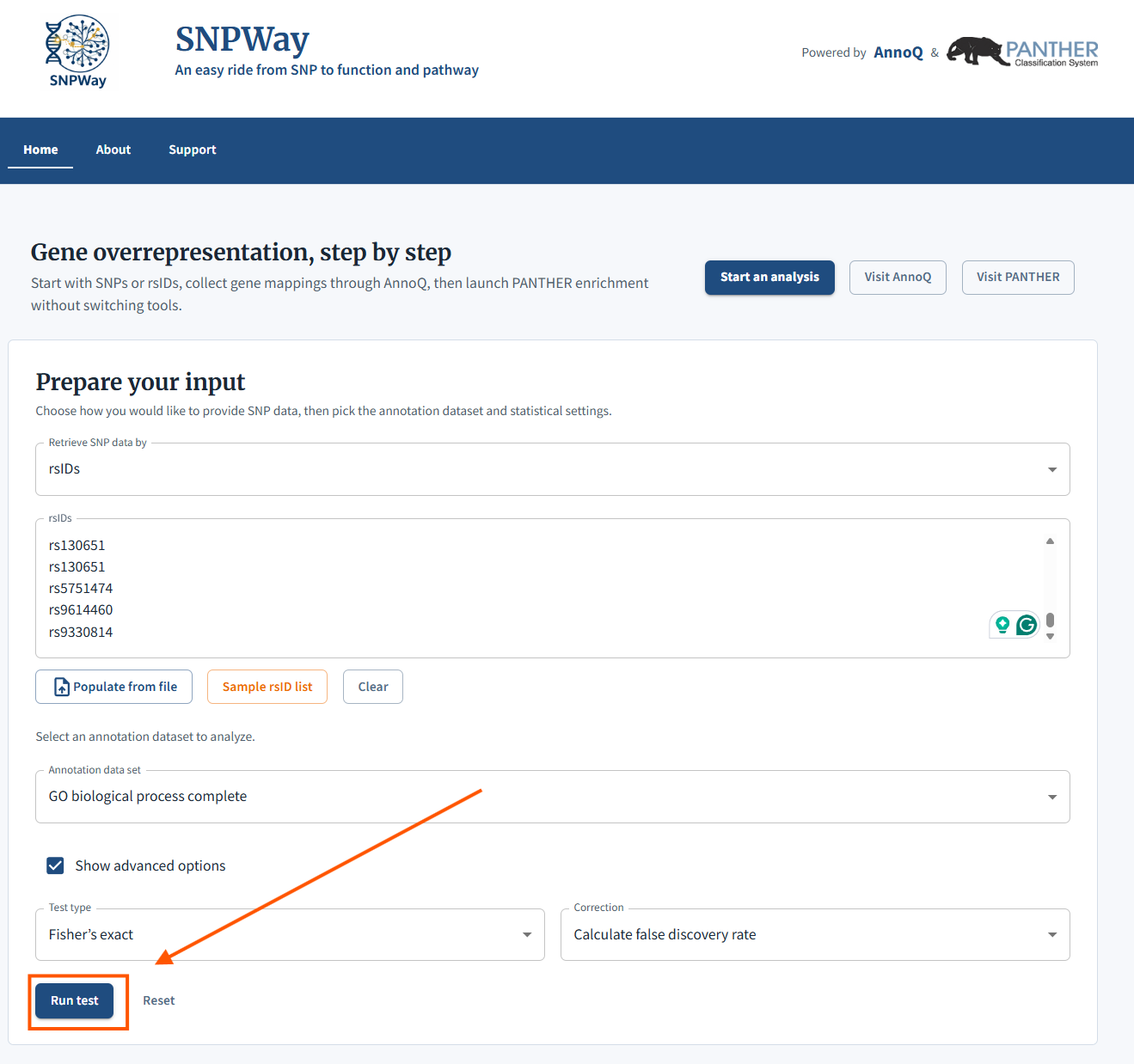

9. A Result window will appear as show below. If you’d like to see more in detail information, you can launch the PANTHER website

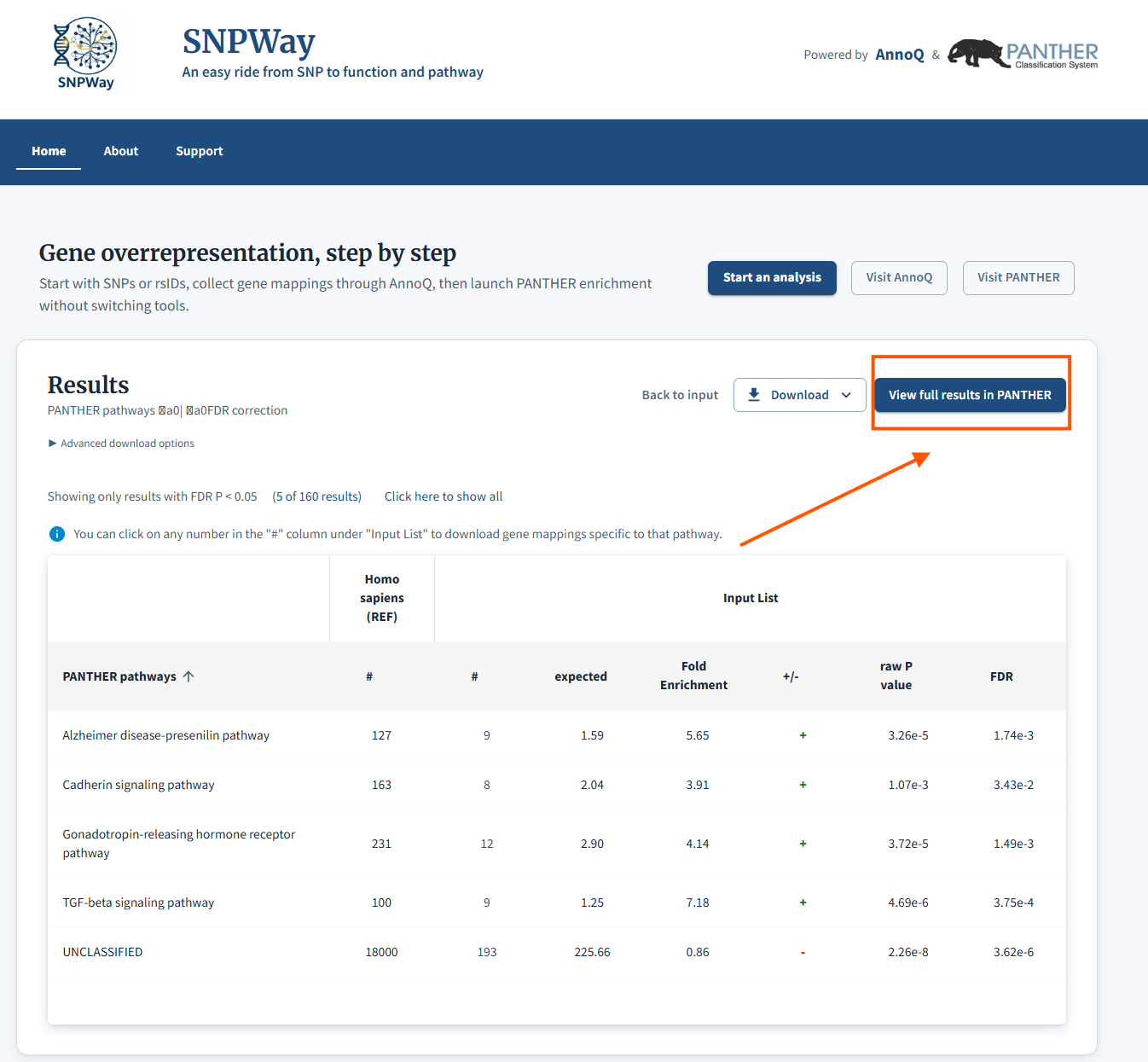
